# Artificial initiation codons and engineered initiator tRNAs enable N-terminal noncanonical amino acid incorporation in intact cell-free translation systems

**DOI:** 10.64898/2026.05.24.725928

**Authors:** Haruyuki Furukawa, Yasunori Okamoto, Naohiro Terasaka

## Abstract

Noncanonical amino acid (ncAA) incorporation at the protein *N*-terminus provides a powerful strategy for installing defined chemical handles while minimizing perturbation of internal protein sequences. However, highly efficient initiation-based ncAA incorporation systems suppress the native methionine pathway by removing methionine or methionyl-tRNA synthetase, limiting their use for proteins containing internal methionine residues. Here, we developed an orthogonal initiation system for selective *N*-terminal ncAA incorporation into proteins in intact cell-free translation systems. We systematically profiled background initiation from all 64 codons in reconstituted translation systems and identified low-background artificial initiation codons. Engineered initiator tRNAs, termed tRNA^IniTx^, were then designed to decode selected codons and support ncAA-dependent initiation. The optimized CAC/tRNA^IniTx04^_GUG_ pair enabled efficient *N*-terminal incorporation of *N*-biotinyl-L-phenylalanine without removing methionine or methionyl-tRNA synthetase, reaching over 90% incorporation. The system was further extended to *p*-azido-L-phenylalanine and to an *Escherichia coli* extract-based cell-free translation system. Finally, *N*-terminally biotinylated proteins were directly immobilized on streptavidin biosensors for purification-free biolayer interferometry analysis of computationally designed Brd4^BD2^ binders. This work establishes a codon-guided orthogonal initiation strategy for *N*-terminal protein functionalization while preserving the native methionine translation pathway.

## INTRODUCTION

Noncanonical amino acid (ncAA) incorporation expands the chemical functionality of proteins beyond the repertoire provided by the 20 canonical amino acids. By installing ncAAs at defined positions, proteins can be equipped with chemical handles for labeling, conjugation, or interaction analysis, thereby enabling applications that are difficult to achieve by conventional mutagenesis alone. Cellular genetic code expansion systems are well suited for large-scale production of ncAA-containing proteins^1,2^ and intracellular functional analyses^3–5^. However, the chemical diversity and number of ncAAs that can be incorporated in cells remain constrained by cellular viability and compatibility with endogenous translation machinery^6^. In contrast, *in vitro* translation systems^7^ allow direct manipulation of translation components regardless of cellular viability, which enables rapid synthesis of diverse ncAA-containing proteins.

*In vitro* ncAA incorporation can be achieved through either translation elongation or initiation. Amber suppression^8^ and four-base codon translation systems^9^ are powerful for introducing ncAAs at internal positions of proteins. In contrast, initiation-based ncAA incorporation provides a distinct route to *N*-terminal protein functionalization. Because the initiating amino acid is positioned at the ribosomal P site and its α-amino group is not required as the nucleophile for peptide-bond formation, translation initiation can accommodate *N*-modified and structurally diverse amino acid derivatives^10^. This feature makes initiation-based incorporation especially useful for installing a single chemical handle at the *N*-terminus while leaving the internal protein sequence unaltered. Such *N*-terminal functionalization is attractive for cell-free workflows in which protein synthesis can be directly coupled to downstream immobilization, conjugation, or interaction assays.

*N*-terminal ncAA incorporation has been extensively developed in genetic-code-reprogrammed *in vitro* translation systems^11^. In these systems, methionine and/or methionyl-tRNA synthetase (MetRS) are removed, allowing engineered initiator tRNAs charged with ncAAs to be efficiently utilized at the initiation step. These systems have been widely used for ribosomal synthesis of nonstandard peptides such as cyclic peptides^12^ and peptides containing fluorescein^13^, long-chain fatty acids^14^ and exotic amino acids^15^. These genetic-code-reprogrammed studies have also shown that translation initiation is not strictly limited to the AUG codon. Reprogrammed “dual-sense codon” systems demonstrated that selected codons can assign distinct amino acids at both initiation and elongation, thereby expanding the repertoire of initiators and elongators that can be used simultaneously^16^. EF-P-responsive artificial initiator tRNAs further enabled efficient *N*-terminal incorporation of difficult substrates, including D-amino acids, β-amino acids, and γ-amino acids, and showed that non-AUG codon–anticodon pairs such as AAG/CUU and GUA/UAC can support efficient initiation under optimized conditions^15^. More recently, engineered initiator tRNAs carrying diverse anticodons were systematically evaluated in a genetically reprogrammed in vitro translation system, demonstrating that multiple non-AUG codons can function as initiation codons and that tRNA folding strongly influences initiation efficiency^17^.

Despite these advances, genetic-code-reprogrammed systems are generally optimized for peptide synthesis under conditions in which the native methionine translation pathway, including endogenous fMet-tRNA^fMet^ formation, is suppressed. As a result, they are not readily applicable to the synthesis of proteins containing internal methionine residues. For such applications, ncAA incorporation must be confined to an artificial initiation event that is functionally orthogonal to endogenous fMet-tRNA^fMet^-dependent initiation, while preserving the native 20-amino-acid translation system during elongation. Such orthogonal initiation has previously been explored using the amber codon (UAG) as an artificial initiation codon. Early studies showed that replacing the AUG initiation codon with UAG, together with an engineered initiator tRNA bearing a cognate CUA anticodon, can redirect translation initiation from AUG to an amber codon^18^. This principle was later applied to cell-free *N*-terminal protein labeling, in which chemically aminoacylated amber initiator suppressor tRNAs introduced fluorescent or biotinylated amino acids at the *N*-terminus of proteins from templates bearing a UAG initiation codon^19,20^.

Although UAG-based artificial initiation established the feasibility of redirecting translation initiation to a noncanonical initiation codon, it remains unclear whether other codons can serve as artificial initiation codons in intact cell-free translation systems that preserve the native methionine pathway. This question is important because endogenous fMet-tRNA^fMet^ remains present under intact translation conditions and can initiate translation from non-AUG codons to varying degrees, generating background protein synthesis in the absence of the desired ncAA-charged initiator tRNA. Indeed, systematic measurements in *Escherichia coli* cells have shown that many non-AUG codons can initiate translation^21^. Thus, designing orthogonal artificial initiation systems requires identifying codon–initiator tRNA pairs that both minimize endogenous background initiation and support efficient ncAA-dependent initiation while maintaining internal methionine incorporation during elongation.

Here, we developed a methionine-compatible orthogonal initiation system for selective *N*-terminal ncAA incorporation in *E. coli* based intact cell-free translation systems (Fig. 1). Building on previous UAG-initiation and engineered initiator tRNA studies, we first systematically profiled endogenous background initiation from all 64 codons in reconstituted cell-free translation systems. We then engineered corresponding initiator tRNA, termed tRNA^IniTx^, to enable efficient ncAA-dependent initiation at selected low-background artificial initiation codons. Using this strategy, we achieved efficient *N*-terminal incorporation of *N*-biotinyl-L-phenylalanine (BioPhe) without removing methionine or MetRS. We further demonstrate compatibility with *p*-azido-L-phenylalanine, T-boxzyme-mediated aminoacylation^22^, and *E. coli* extract-based cell-free translation system. Finally, we show that *N*-terminal biotinylation enables direct immobilization of cell-free synthesized proteins for purification-free biolayer interferometry analysis, including the evaluation of computationally designed protein binders. Together, this work establishes a codon-guided strategy for *N*-terminal protein functionalization while preserving the native methionine translation pathway.

**Figure 1.**
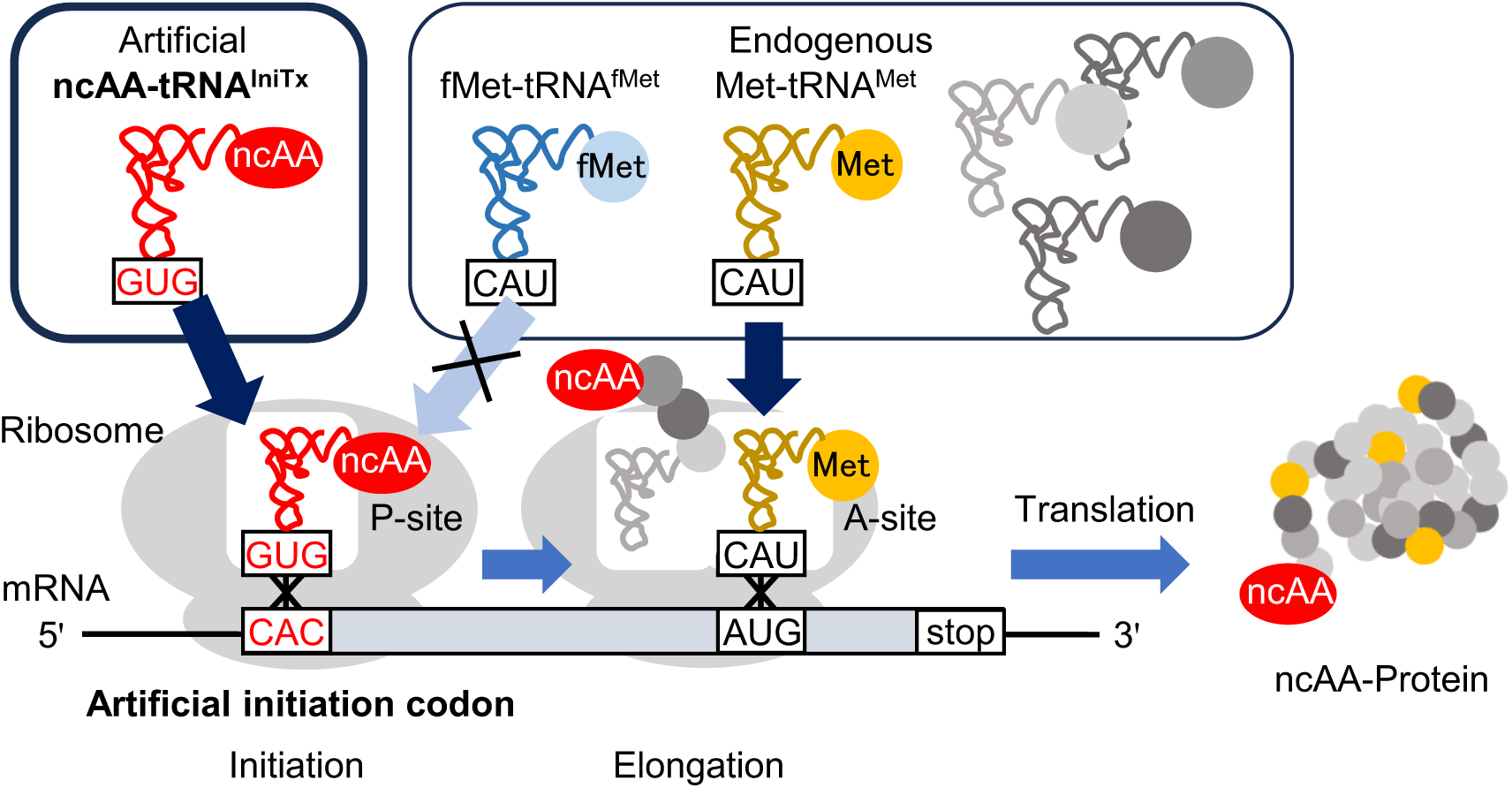
Schematic illustration of orthogonal translation initiation for selective N-terminal ncAA incorporation. An artificial initiation codon poorly recognized by endogenous fMet-tRNA^fMet^ is selectively decoded by ncAA-tRNA^IniTx^. Endogenous Met-tRNA^Met^ is utilized for methionine incorporation at elongation AUG codons.

## RESULTS

### Background initiation from all 64 codons in reconstituted cell-free translation systems

To identify artificial initiation codons that minimize undesired initiation, we systematically analyzed translation output from all 64 initiation codons using an mNeonGreen (mNG) reporter in which all sequences except the initiation codon were identical. Translation reactions were performed using PUREfrex 1.0 and PUREfrex 2.1, two differently optimized commercially available reconstituted cell-free translation systems, to evaluate whether translation-system composition and reaction conditions affect non-AUG initiation. Reactions were performed in the absence of engineered initiator tRNAs and ncAA-charged tRNAs. Under these conditions, fluorescence output from non-AUG templates was used as a measure of background initiation by endogenous translation components.

Translation output varied substantially among initiation codons in both reconstituted cell-free translation systems (Fig. 2), demonstrating that codon identity strongly influences background initiation under intact cell-free translation conditions. Overall, AU-rich codons tended to exhibit higher background initiation activity, whereas GC-rich codons generally showed lower activity. In particular, many highly active codons contained U at the second nucleotide position, partially consistent with previous measurement of translation initiation from all 64 initiation codons in *E. coli* cells^21^. The detailed profiles of initiation differed between PUREfrex 1.0 and PUREfrex 2.1. Translation output from many non-AUG codons in PUREfrex 2.1 was reduced, suggesting higher stringency against non-AUG initiation. Importantly, several codons exhibited lower background initiation activity than UAG codon commonly used in previous UAG-based orthogonal initiation systems^18–20^. Among these candidates, CAC, UCC and CCC consistently exhibited low background initiation activity in both PUREfrex systems. These codons were therefore selected as candidate low-background artificial initiation codons for subsequent evaluation with engineered initiator tRNAs.

**Figure 2.**
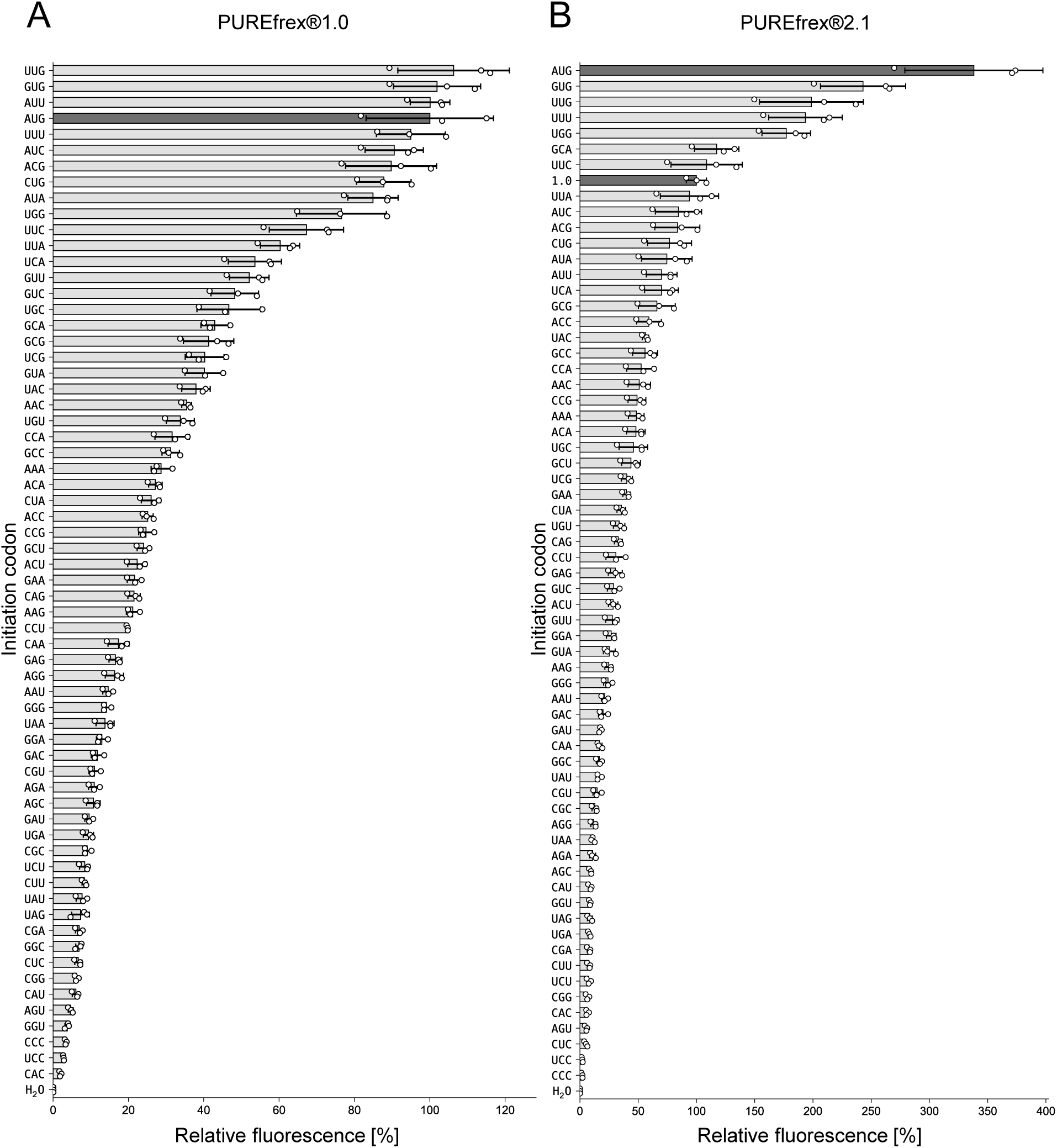
Translation initiation efficiency from 64 codons in PURE systems. DNA templates encoding mNG bearing various initiation codons or water were added to PUREfrex 1.0 **(A)** or PUREfrex 2.1 (**B**). Relative fluorescence intensity was normalized to that of mNG expressed from DNA template bearing AUG initiation codon in PUREfrex1.0 was shown. Bars represent the mean ± s.d. of three independent reactions (n = 3). Initiation codons are ordered according to decreasing translation output.

### Engineering of initiator tRNA for selective N-terminal incorporation of BioPhe

To enable ncAA-dependent initiation at the selected low-background codons, we designed a series of engineered initiator tRNAs, termed tRNA^IniTx^, based on the previously reported tRNA^IniP^ scaffold^15^. tRNA^IniP^ is a chimeric initiator tRNA designed to improve *N*-terminal incorporation of difficult substrates by combining initiator tRNA features with tRNA^Pro^ elements recognized by elongation factor P (EF-P). In the present study, the anticodon of tRNA^IniP^ was replaced with cognate sequences to the selected artificial initiation codons. In addition, ΔC17/C17a mutations were introduced into the D-arm to enable recognition by the tRNA-recognizing aminoacylation ribozyme T-boxzyme^22^. Because anticodon replacement and D-arm engineering can perturb tRNA folding, the secondary structures of the designed tRNAs were evaluated using RNAfold^23^ (ViennaRNA Package 2.0). Based on these predictions, we additionally introduced a previously reported tRNA^Ini^/tRNA^Pro^ chimeric motif^15^ to restore tRNA-like structures^17^. The resulting five tRNA^IniTx^ variants are summarized in Supplementary Fig. S1.

We next evaluated whether the tRNA^IniTx^ variants could support *N*-terminal ncAA incorporation using *N*-biotinyl-L-phenylalanine (BioPhe) as a model ncAA. BioPhe-CME (Fig. 3A), the cyanomethyl ester-activated form of BioPhe, is a substrate for aminoacylation ribozymes^11,22^. In addition, BioPhe incorporation can be detected by streptavidin (SAv) mediated electrophoresis mobility shift assay. Translation products were analyzed by non-boiled SDS-PAGE in the absence or presence of SAv, followed by fluorescence detection of mNG^24^. We first examined BioPhe incorporation using CAC initiation codon and tRNA^IniTx04^_GUG_ pair in PUREfrex 1.0 without EF-P (Fig. 3B). In the absence of tRNA^IniTx04^_GUG_, only low translation output was observed. Addition of non-acylated tRNA^IniTx04^ modestly increased translation output, whereas addition of BioPhe-tRNA^IniTx04^_GUG_ prepared using flexizyme (eFx)^11^ markedly increased production of mNG. SAv-dependent mobility shift was observed only in the presence of BioPhe-tRNA^IniTx04^_GUG_, indicating tRNA-dependent incorporation of BioPhe into the translated protein.

**Figure 3.**
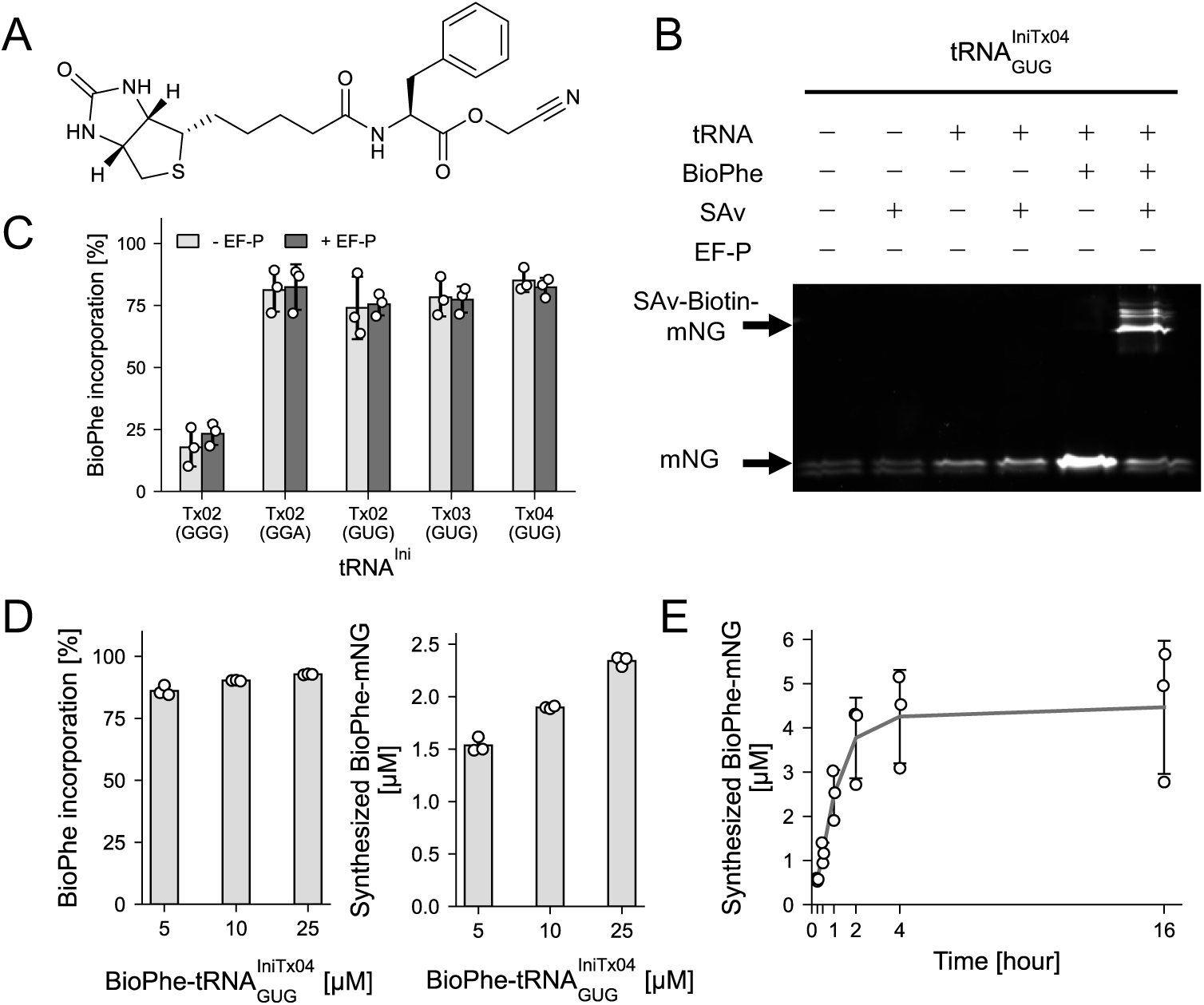
Optimization of orthogonal initiation using selected artificial initiation codons and tRNA^IniTx^ variants for BioPhe incorporation. (**A**) Chemical structure of BioPhe-CME. (**B**) Detection of BioPhe incorporation using tRNA^IniTx04^_GUG_ into mNG by SAv gel-shift assay in non-boiled SDS–PAGE. (**C**) BioPhe incorporation efficiencies using five tRNA^IniTx^ variants cognate to selected artificial initiation codons. (**D**) Effect of BioPhe-tRNA^IniTx04^_GUG_ concentration on incorporation efficiency (left panel) and absolute yield of BioPhe-incorporated mNG (right panel). (**E**) Time-course analysis of BioPhe incorporation mediated by tRNA^IniTx04^_GUG_. Quantitative data (panels C, D and E) represent the mean ± s.d. of three independent translation reactions (n = 3).

We then compared BioPhe incorporation among the five tRNA^IniTx^ variants corresponding to the selected artificial initiation codons (Supplementary Figs. S2 and S3, and Fig. 3C). BioPhe incorporation efficiency was calculated from the fraction of the SAv-shifted band relative to total mNG fluorescence signal. All tested tRNA^IniTx^ variants supported tRNA-dependent BioPhe incorporation, with tRNA^IniTx04^_GUG_ showing the highest incorporation efficiency. Addition of EF-P did not substantially improve BioPhe incorporation under the tested conditions. Using tRNA^IniTx04^, we optimized the concentration of BioPhe-tRNA^IniTx04^ (Fig. 3D). High incorporation efficiencies of approximately 90% were observed across all conditions, while protein synthesis yields ranged from approximately 1.5 to 2.5 μM. A time-course analysis using 10 μM BioPhe-tRNA^IniTx04^_GUG_ showed that BioPhe-mNG production reached a plateau within approximately 4 h and remained largely stable up to 16 h (Fig. 3E). Based on these results, the CAC/tRNA^IniTx04^_GUG_ pair was selected for subsequent experiments.

We also evaluated BioPhe incorporation using PUREfrex 2.1. Although tRNA^IniTx^-dependent SAv-shifted bands were detected, the maximum incorporation efficiency was limited to approximately 15%, substantially lower than that observed in PUREfrex 1.0 (Supplementary Figs. S4-S6). These results indicate that tRNA^IniTx^-dependent artificial initiation can occur in both PUREfrex systems, but that PUREfrex 1.0 provides a more favorable balance between low-background initiation and efficient tRNA^IniTx^-dependent incorporation.

### Extending the orthogonal initiation platform to T-boxzyme aminoacylation, an azido ncAA, and extract-based translation

To evaluate the versatility of the developed orthogonal initiation platform, we examined whether the CAC/tRNA^IniTx04^_GUG_ pair could be extended beyond the initial BioPhe/eFx/PUREfrex 1.0 system. We tested three extensions in a stepwise manner: replacement of eFx with the tRNA-recognizing aminoacylation ribozyme T-boxzyme, replacement of BioPhe with *p*-azido-L-phenylalanine (AzPhe), and replacement of the reconstituted PURE system with an *E. coli* extract-based cell-free translation system.

We first tested T-boxzyme-mediated aminoacylation of tRNA^IniTx^. Because tRNA^IniTx^ variants were designed to contain D-arm modifications for T-boxzyme recognition, this experiment directly evaluated whether tRNA^IniTx^ design was compatible with a tRNA-recognizing aminoacylation ribozyme. BioPhe-charged tRNA^IniTx^ prepared using T-boxzyme (Tx2.1) supported SAv-dependent band shifting in PUREfrex 1.0, confirming BioPhe incorporation into mNG (Fig. S7A). Screening and optimization identified tRNA^IniTx04^_GUG_ aminoacylated using 20 μM Tx2.1 and 20 μM tRNA as the best condition (Supplementary Fig. S7B, C), yielding an apparent BioPhe incorporation efficiency of 93.8 ± 1.1% and BioPhe-mNG production of 2.6 ± 0.6 μM. These results demonstrate that the CAC/tRNA^IniTx04^_GUG_ pair is compatible not only with eFx-mediated aminoacylation, but also with Tx2.1-mediated aminoacylation.

We next asked whether the optimized artificial initiation system could incorporate an ncAA other than BioPhe. As a clickable ncAA, we tested *p*-azido-L-phenylalanine^25^. AzPhe activated with CME (AzPhe-CME, Supplementary Fig. S8A) was used for aminoacylation by eFx and further *N*-acetylated according to a previous report^15^. The resulting Ac-AzPhe-tRNA^IniTx04^_GUG_ was then used for translation of a CAC-start mNG DNA template in PUREfrex 1.0. To detect AzPhe incorporation, the translated protein was biotinylated through copper-free click chemistry using sulfo-DBCO-biotin, desalted to remove excess reagent, and analyzed by SAv gel-shift assay (Supplementary Fig. S8B). SAv-dependent shifted bands were detected only when Ac-AzPhe-charged tRNA^IniTx04^_GUG_ was supplied, confirming ncAA-dependent incorporation of Ac-AzPhe (Supplementary Fig. S8C). Addition of 50 μM Ac-AzPhe-charged tRNA^IniTx04^_GUG_ to PUREfrex 1.0 yielded an incorporation efficiency of 67.2 ± 9.7% and a protein yield of 1.9 ± 0.3 μM (Supplementary Fig. S8D).

Finally, we examined whether the system could be applied to an *E. coli* extract-based cell-free translation system. BioPhe-tRNA^IniTx04^_GUG_ was added together with a CAC start mNG DNA template. As observed in PUREfrex 1.0, SAv-dependent shifted bands were detected only in the presence of BioPhe-tRNA^IniTx04^_GUG_, indicating successful *N*-terminal BioPhe incorporation in the crude extract-based system (Supplementary Fig. S9A). Optimization of BioPhe-tRNA concentration showed that 5 and 10 μM conditions yielded incorporation efficiencies of approximately 90% with production of approximately 0.07 μM (Supplementary Fig. S9B). These results demonstrate that the orthogonal initiation platform is not limited to reconstituted translation systems and can be extended to crude extract-based cell-free translation systems.

### Purification-free BLI analysis of de novo designed protein binders

As an application of the developed BioPhe incorporation system, we performed purification-free biolayer interferometry (BLI) analysis of protein binders, inspired by FASTIA (Fast Affinity Screening Technology for Interaction Analysis)^26^. We designed 14 mini-protein binders targeting human Brd4^BD2^ using the AI-based binder design pipeline BindCraft^27^. Brd4^BD2^ is an attractive therapeutic target for various diseases^28^. *N*-terminally BioPhe-labeled mNG-fused binders were synthesized using PUREfrex 1.0 and immobilized directly onto SAv-coated biosensors. The analyte, mScarlet-fused Brd4^BD2^, was synthesized using PUREfrex 2.1. Both ligands and analytes were quantified by fluorescence and used directly for BLI measurements without purification (Fig. 4A).

**Figure. 4.**
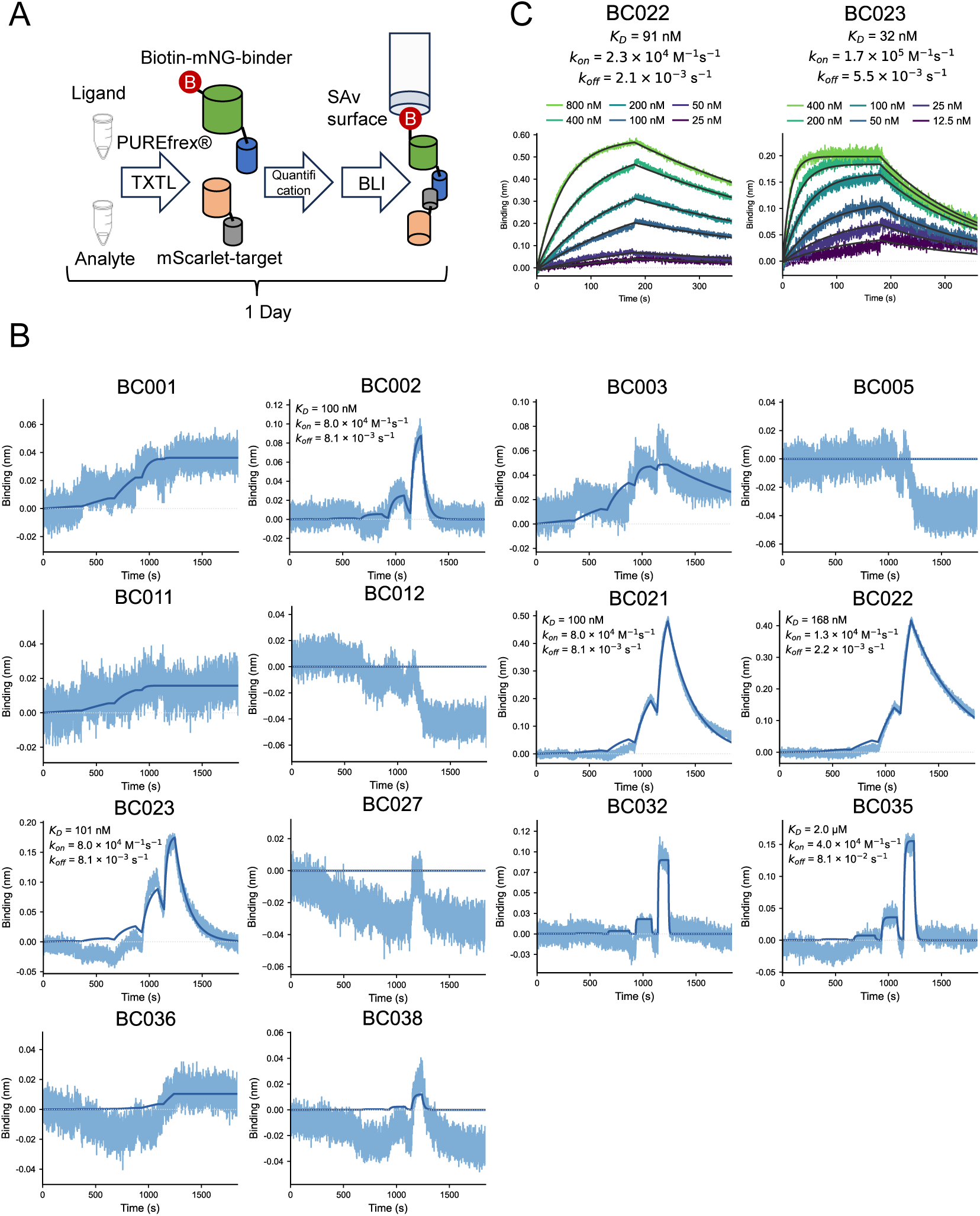
BLI screening of de novo designed binders using cell-free expressed biotinylated proteins. (**A**) Schematic workflow of BLI analysis. BioPhe-mNG-fused Brd4^BD2^-targeting binders were expressed by cell-free translation, quantified by fluorescence, and directly immobilized on SAv biosensors. (**B**) Screening of binders by single-cycle kinetics (SCK) BLI measurements. The blue lines represent the reference-subtracted data, and the black lines represent the fitting curves. (**C**) Detailed kinetic analysis of selected binders by multi-cycle kinetics (MCK) BLI measurements. The colored lines represent the reference-subtracted data, and the black lines represent the fitting curves.

We first confirmed expression and BioPhe incorporation for the 14 mNG-fused binders. CAC-start DNA templates were used in PUREfrex 1.0 in the presence of 5 μM BioPhe-tRNA^IniTx04^_GUG_. Non-boiled SDS–PAGE analysis in the presence of SAv confirmed shifted bands for all binders, indicating successful BioPhe incorporation (Supplementary Fig. S10). We then performed single-cycle kinetic BLI screening. Among the 14 tested binders, four exhibited apparent *K*_D_ values in the range of approximately 100–200 nM (Fig. 4B). Representative binders BC022 and BC023 were further analyzed by multi-cycle kinetic measurements. BC022 exhibited a slower *k*_off_ value than BC023, demonstrating that the platform can distinguish binders with different kinetic properties directly from cell-free translation reactions (Fig. 4C). These results show that *N*-terminal BioPhe incorporation can directly couple cell-free synthesis of designed proteins to purification-free interaction analysis.

## DISCUSSION

In this study, we established an orthogonal translation initiation system that enables selective *N*-terminal incorporation of noncanonical amino acids (ncAAs) in intact cell-free translation systems. Conventional strategies for *N*-terminal ncAA incorporation generally suppress competition from endogenous initiator tRNAs by removing Met and/or MetRS from reconstituted translation systems^12,15,17,29^. Although such genetic-code-reprogrammed systems are powerful for peptide synthesis, they are poorly suited for the synthesis of proteins containing internal methionine residues. In contrast, our strategy preserves the native methionine translation pathway during elongation while redirecting initiation to artificial initiation codons. This design enables *N*-terminal ncAA incorporation without Met/MetRS depletion and therefore expands the applicability of initiation-based ncAA incorporation from specialized peptide synthesis toward methionine-containing proteins and cell-free protein workflows.

To identify codons suitable for orthogonal initiation, we systematically profiled background initiation from all 64 codons in intact reconstituted translation systems. Previous UAG-based artificial initiation studies established that translation initiation can be redirected to an artificial initiation codon, but they did not define whether other codons could provide lower background initiation in intact cell-free systems^18–20^. Our 64-codon analysis showed that background initiation strongly depends on codon identity and that several codons exhibit lower background activity than the UAG. The identification of CAC, UCC, and CCC as low-background candidates provided the basis for engineering codon–tRNA pairs that combine background suppression with efficient ncAA-dependent initiation.

The comparison between PUREfrex 1.0 and PUREfrex 2.1 further highlights the importance of translation-system context in orthogonal initiation design. PUREfrex 2.1 showed reduced translation output from many non-AUG codons, suggesting higher stringency against non-AUG initiation. However, this increased stringency also coincided with reduced tRNA^IniTx^-dependent BioPhe incorporation. When adding tRNA^IniTx^ variants without aminoacylation to PUREfrex 2.1, background initiation was significantly enhanced (Supplementary Figs. S3 and S4). This observation suggests that components or factor stoichiometries in PUREfrex 2.1 may enhance charging or utilization of non-acylated tRNA^IniTx^ variants, although the detailed mechanism remains unclear. In contrast, PUREfrex 1.0 provided a more favorable balance for the present platform. Low-background codons could suppress endogenous initiation, while tRNA^IniTx^ variants could still support efficient ncAA-dependent initiation. These results suggest that the optimal translation system for orthogonal initiation is not necessarily the system with the lowest non-AUG initiation activity, but rather the system that best balances endogenous background suppression and engineered tRNA-dependent initiation.

The performance of the engineered initiator tRNAs likely reflects multiple interdependent factors. The tRNA^IniTx^ variants were designed by combining anticodon replacement, D-arm engineering for T-boxzyme recognition, and secondary structure compensation using a previously reported tRNA^Ini^/tRNA^Pro^ chimeric motif^15,17^. Among the tested variants, tRNA^IniTx04^_GUG_ and CAC initiation codon exhibited the highest BioPhe incorporation efficiency. The performance of tRNA^IniTx^ variants is likely governed by multiple factors, including codon–anticodon interactions, initiator tRNA identity, structural stability, aminoacylation efficiency, IF2 recognition, ribosome accommodation, and competition with endogenous initiation. Thus, although the present study identifies a highly efficient CAC/tRNA^IniTx04^_GUG_ pair, further systematic mutational analysis will be useful for defining general design rules for artificial initiator tRNAs.

Several mechanistic aspects remain to be clarified. In the BioPhe gel-shift analysis, CAC-start mNG showed a doublet band pattern under non-boiled SDS-PAGE conditions (Supplementary Figs. S2-S5). The relative intensities of these bands changed upon addition of tRNA^IniTx^ and BioPhe-charged tRNA^IniTx^, suggesting that they may reflect heterogeneous initiation products or altered electrophoretic behavior caused by different *N*-terminal compositions. Direct mass spectrometric characterization would be useful for defining the molecular identity and positional fidelity of the *N*-terminally modified products. In addition, although tRNA^IniTx^ variants were derived from an EF-P-responsive tRNA^IniP^ scaffold, addition of EF-P did not substantially improve BioPhe incorporation under the tested conditions. This may reflect partial disruption of the EF-P recognition motif by the ΔC17/C17a mutations, or the relatively high compatibility of BioPhe with initiation in this system.

The developed platform was compatible with multiple extensions beyond the initial BioPhe/eFx/PUREfrex 1.0 condition. First, BioPhe incorporation using T-boxzyme-mediated aminoacylation demonstrated that the tRNA^IniTx^ design is compatible with a tRNA-recognizing aminoacylation ribozyme. This is notable because T-boxzyme can aminoacylate cognate tRNA in situ in customized PURE system^22^. Further optimization and tRNA/T-boxzyme engineering may enable *in situ* ncAA incorporation at *N*-terminus of proteins. Second, incorporation of Ac-AzPhe showed that the system can introduce a clickable *N*-terminal residue, enabling downstream conjugation through copper-free click chemistry. Third, BioPhe incorporation in an *E. coli* extract-based cell-free translation system indicated that the CAC/tRNA^IniTx04^_GUG_ strategy is not limited to purified reconstituted systems. Together, these results demonstrate that the platform can be adapted across aminoacylation chemistries, ncAA substrates, and cell-free translation formats.

As a proof-of-concept application, we used the orthogonal initiation system to directly immobilize cell-free synthesized BioPhe-containing proteins onto SAv-coated biosensors for purification-free BLI analysis. Because biotin-SAv interaction has extremely high affinity, our platform enables stable immobilization avoiding baseline drift during BLI measurement, which is advantageous to get reliable data in the initial binder screening steps. The successful screening of computationally designed Brd4^BD2^ binders and subsequent kinetic characterization of selected binders show that the method can support rapid evaluation of designed protein variants. Such workflows should be particularly useful for protein engineering and AI-assisted protein design such as RFdiffusion^30^, BindCraft^27^, and BoltzGen^31^, where many methionine-containing variants may need to be synthesized and functionally evaluated in parallel.

Taken together, this study establishes an orthogonal translation initiation system for N-terminal ncAA incorporation in intact cell-free systems, which is compatible with the native methionine translation machinery. This approach is useful for cell-free workflows that require the expression of methionine-containing protein and direct coupling of protein synthesis to downstream immobilization, conjugation, or interaction analysis.

## MATERIALS AND METHODS

Polymerase chain reaction (PCR) and primer extension or DNA fragment extension reactions were performed using KOD One PCR Master Mix (KMM-201, TOYOBO). DNA oligonucleotides were purchased from Integrated DNA Technologies or Eurofins Genomics. Gene fragments were obtained from Integrated DNA Technologies or GenScript.

### Chemical synthesis of amino acid substrates

The non-canonical amino acid substrates were synthesized following the previously reported methods^32,33^ with minor modifications as described below.

Biotin-Phe: L-Phenylalanine (435 mg, 2.64 mmol) was suspended in dimethylformamide (DMF, 7.5 mL). To this suspension, a suspension of biotin N-hydroxysuccinimide ester (900 mg, 2.64 mmol) in DMF (7.5 mL) was added, followed by the addition of NaHCO₃ (664 mg, 7.91 mmol). The resulting mixture was stirred at 50 °C for 8 h. The reaction was monitored by TLC (CH₂Cl₂:MeOH:AcOH = 20:80:0.1). The mixture was then acidified to pH 2-3 with 1 N HClaq and stirred at ambient temperature for 30 min. The obtained white precipitate was collected and washed sequentially with diethyl ether, 200 mM HClaq, 10 mM HClaq, and diethyl ether. The resulting white solid (Biotin-Phe) was dried under reduced pressure.

Biotin-Phe-CME: To a solution of Biotin-Phe (350 mg, 893 μmol) in DMF (1.0 mL) were added triethylamine (137 μL, 983 μmol) and chloroacetonitrile (358 μL, 5.65 mmol). The reaction mixture was stirred at ambient temperature overnight. The reaction was monitored by TLC (CH₂Cl₂:MeOH:AcOH = 93:7:0.1). During the reaction, the pH was monitored and aliquots of triethylamine were added to maintain basic conditions (∼pH 9). After addition of ethyl acetate (10 mL), the obtained mixture was washed with 1 N HClaq (10 mL), 20 mM HClaq, saturated aqueous NaHCO₃, and brine. The organic layer was dried over Na₂SO₄, filtered, and concentrated under reduced pressure.

Boc-4-Azido-Phe-CME: Boc-4-azido-L-phenylalanine (800 mg, 2.61 mmol) was dissolved in DMF (1.5 mL). To the solution were added triethylamine (400 μL, 2.87 mmol) and chloroacetonitrile (1.08 mL, 16.98 mmol). The reaction mixture was stirred at ambient temperature overnight. The reaction was monitored by TLC (cyclohexane/ethyl acetate/triethylamine = 50:50:0.1). During the reaction, the pH was monitored and aliquots of triethylamine were added to maintain basic conditions (∼pH 9). The mixture was diluted with diethyl ether (10 mL) and washed with 1 N HClaq (10 mL), 20 mM HClaq, saturated aqueous NaHCO₃, and brine. The organic layer was dried over Na₂SO₄, filtered, and concentrated under reduced pressure. The crude product was purified by silica gel column chromatography (cyclohexane:ethyl acetate:triethylamine = 50:50:0.1), and the solvent was removed under reduced pressure.

4-Azido-Phe-CME: To a solution of Boc-4-azido-Phe-CME (680 mg, 1.97 mmol) in dichloromethane (2.0 mL) was added 4 N HCl in ethyl acetate (6 mL). The reaction mixture was stirred at ambient temperature overnight. The reaction was monitored by TLC (cyclohexane/ethyl acetate = 65:35). The volatiles were removed under reduced pressure. The obtained solid was washed with diethyl ether and dried under reduced pressure.

### Preparation of linear DNA templates for *in vitro* transcription and translation

Linear DNA templates for *in vitro* transcription and translation were prepared by PCR using plasmids as templates and appropriate primers. PCR products were purified using the FastGene Gel/PCR Extraction Kit (FG-91302, NIPPON Genetics) and quantified with the Qubit dsDNA Quantification Assay Kit (Q32854, Thermo Fisher Scientific). DNA oligonucleotide sequences, amino acid sequences of proteins, and nucleotide sequences of DNA templates are provided in Supplementary Tables S1–S7.

DNA templates of mNG with artificial initiation codons were generated by PCR using mutagenic primers. DNA templates for T7pro-CAC-mNG-binders, T7pro-mScarlet-Brd4^BD2^, and T-boxzymes were generated by overlap extension of the corresponding DNA fragments, followed by PCR amplification. For tRNA templates, the primer corresponding to the 3′ end contained 2′-O-methyl modifications at its two 5′-terminal nucleotides to suppress non-templated nucleotides addition by T7 RNA polymerase^34^. Detailed procedures are described in the Supplementary Methods.

### In vitro transcription and purification of RNAs

eFx was prepared according to previously described methods^15^.

DNA templates for tRNAs and T-boxzymes were purified by phenol–chloroform extraction, ethanol precipitation, and a 70% ethanol wash. Transcription reactions for tRNAs were carried out overnight at 37 °C in buffer containing 40 mM Tris–HCl (pH 8.0), 22.5 mM MgCl₂, 1 mM dithiothreitol (DTT), 1 mM spermidine, 0.01% Triton X-100, 3.75 mM nucleoside triphosphate (NTP) mix, 5 mM guanosine monophosphate (GMP), and 0.12 μM in-house expressed T7 RNA polymerase. Transcription reactions for T-boxzymes were carried out overnight at 37 °C in buffer containing 40 mM Tris–HCl (pH 8.0), 30 mM MgCl₂, 1 mM DTT, 1 mM spermidine, 0.01% Triton X-100, 5 mM NTP mix and 0.12 μM in-house expressed T7 RNA polymerase.

The transcription reactions were treated with RQ1 DNase (Promega) at 37 °C for 0.5–1 h. The resulting tRNAs and T-boxzymes were purified by denaturing polyacrylamide gel electrophoresis on 8% (tRNAs) or 6% (T-boxzymes) gels containing 8 M urea and 1×TBE. RNAs were excised from the gels, eluted, precipitated with isopropanol, washed with 70% ethanol, and dissolved in water.

### Preparation of aminoacyl-tRNAs

Preparation of BioPhe-tRNA and acetylated AzPhe-tRNA using eFx was performed based on a previously reported method^12^ with minor modifications, as described below. A mixture containing 83 mM HEPES-KOH (pH 7.5), 42 µM eFx, and 42 µM tRNA variants was incubated at 95°C for 2 min, followed by cooling at room temperature for 5 min. MgCl₂ and amino acid substrates were then added to initiate the aminoacylation reaction. The final reaction mixture contained 50 mM HEPES-KOH (pH 7.5), 25 µM eFx, 25 µM tRNA variants, 600 mM MgCl₂, and 5 mM BioPhe or AzPhe, and was incubated on ice for 1–3 h. Following the reaction, tRNAs were precipitated by the addition of 4 volumes of 0.3 M sodium acetate (pH 5.2) and 10 volumes of ethanol. For BioPhe-tRNA, the pellet was rinsed once with 70% ethanol containing 0.1 M sodium acetate (pH 5.2). For AzPhe-tRNA, the pellet was rinsed twice with the same solution. The pellets were air-dried and dissolved in 1 mM sodium acetate (pH 5.2) prior to use in in vitro translation. Acetylation of AzPhe-tRNA was performed as described previously^15^. The resulting N-acetyl-aminoacyl-tRNAs were washed twice with 70% ethanol containing 0.1 M sodium acetate (pH 5.2) and dissolved in 1 mM sodium acetate (pH 5.2).

Aminoacylation of tRNA^IniTx^ using a T-boxzyme was performed based on a previously reported method^22^ with minor modifications. A mixture of T-boxzyme and tRNA variants was incubated at 95°C for 2 min, followed by cooling at room temperature for 5 min. Reagents were then added to initiate the aminoacylation reaction. The final reaction mixture contained 50 mM HEPES-KOH (pH 7.5), 10, 5, or 2.5 µM T-boxzyme, 10 or 5 µM tRNA variants, 10 mM MgCl_2_, 500 mM KCl, and 5 mM BioPhe, and was incubated on ice for 1–3 h. Following the reaction, tRNAs were precipitated by adding 4 volumes of 0.3 M sodium acetate (pH 5.2) and 10 volumes of ethanol. The pellet was rinsed with 70% ethanol containing 0.1 M sodium acetate (pH 5.2), air-dried, and dissolved in 1 mM sodium acetate (pH 5.2).

### In vitro translation

In vitro translation reactions were performed using PUREfrex 1.0 or 2.1 (GeneFrontier) according to the manufacturer’s recommended protocol. DNA templates were added at a final concentration of 2 nM, and translation reactions were carried out at 37 °C. For translation reactions involving ncAA incorporation, tRNA charged with the ncAA was additionally supplied to the reaction mixture.

For *E. coli* extract–based translation, the Cell-Free Protein Expression Kit “N Mini” (FUJIFILM, A235-0300) was used according to the manufacturer’s instructions. Translation reactions were performed under the recommended reaction conditions at 30 °C for 90 min.

### Expression and purification of mNG and mScarlet proteins

*E. coli* BL21-Gold(DE3) cells (Agilent Technologies) were transformed with pET-mNG or pRSETb-SKIK-His-mScarlet-I3. Expression and purification of mNG were performed as described previously^35^ and the same procedure was applied for mScarlet.

### Fluorescence Measurement and Quantification of mNG and mScarlet

mNG fluorescence was measured using a Synergy H1 microplate reader (BioTek Instruments). Briefly, 5 µL of PUREfrex 1.0 or 4 µL of PUREfrex 2.1 translation reaction was transferred to a black Corning 384-well low-volume round-bottom microplate, and fluorescence was measured with excitation at 485 nm and emission at 528 nm using top optics. mScarlet fluorescence was measured under similar conditions with excitation at 569 nm and emission at 594 nm using top optics. All measurements were performed as endpoint fluorescence and averaged from 10 reads per well.

For quantification, fluorescence intensities of mNG- and mScarlet-fusion proteins were converted to protein concentrations using standard curves generated from purified mNG and mScarlet proteins, respectively, measured under identical conditions.

### SAv-mediated gel-shift assay under non-denaturing conditions

Briefly, 2.0–2.25 µL of the in vitro translation reaction mixture was mixed with 1 µL of 5 mg/mL SAv. SDS sample buffer was then added, and the samples were subjected to 15% SDS-PAGE without boiling at 180 V for 50 min. Fluorescence signals were detected using a Typhoon FLA 9500 scanner (Cytiva) with the Alexa Fluor 488 detection channel.

Band intensities corresponding to the shifted and unshifted mNG were quantified using ImageQuant after background subtraction. BioPhe incorporation efficiency (%) was calculated using the following equation: shifted band intensity / (shifted band intensity + unshifted band intensity). For absolute quantification of protein amounts, a calibration curve was generated using purified mNG, and protein mass was determined from the band intensities. mScarlet or mScarlet fusion proteins were detected using a Typhoon FLA 9500 scanner (Cytiva) with the Cy3 detection channel.

For Ac-AzPhe-incorporated mNG, 1.25 µL of 5 mM Sulfo-DBCO-biotin was added to 5 µL of the post-translation reaction mixture and incubated at 25 °C for 1 h. Subsequently, 23.75 µL of 1× PBS was added, and the sample was desalted using Zeba™ Spin Desalting Columns (7K MWCO, 0.5 mL; Thermo Fisher Scientific, 89882). Of the 30 µL eluate, 6 µL was mixed with 4 µL of 5 mg/mL SAv and analyzed by SDS–PAGE under non-denaturing conditions.

### BLI-based SCK and MCK analysis

BLI measurements were performed using an Octet RED384 system (Sartorius, Göttingen, Germany). Octet® Streptavidin (SA) Biosensors (18-5019, Sartorius) were hydrated in PBS-T-MS (PBS containing 0.05% (v/v) Tween-20, 0.6 M sucrose, and 20 mM MgCl_2_)^26^ for at least 10 min. BioPhe-mNG-binders were expressed using the PURE system, quantified based on fluorescence, diluted in PBS-T-MS to 50 nM, and immobilized onto the biosensors for 5 min. The sensors were then equilibrated in PBS-T-MS for 10 min.

For SCK measurements, experiments were performed with minor modifications based on a previous report^36^. Analyte concentrations were increased stepwise in PBS-T-MS (0.64, 3.2, 16, 80, and 400 nM), and five association–dissociation cycles were conducted. Association phases were performed for 300, 250, 200, 150, and 100 s, respectively, from the lowest to the highest analyte concentrations. Dissociation phases were set to 60 s for each cycle, except for the final cycle, which was extended to 600 s.

For MCK measurements, seven analyte concentrations (12.5, 25, 50, 100, 200, 400, and 800 nM) were prepared in PBS-T-MS, and each concentration was measured in a single cycle consisting of a 180 s association phase followed by a 180 s dissociation phase.

The obtained sensorgrams were processed using double referencing. A reference sensor prepared using a PUREfrex reaction mixture lacking binder was used to subtract nonspecific analyte binding to the sensor surface. In addition, responses from mScarlet alone to each immobilized sensor were subtracted to correct for baseline drift. SCK data were analyzed using the SpyBLI cell-free pipeline as described in a previous study^36^. MCK data were processed and fitted using a publicly available Python analysis pipeline (available at: https://github.com/chan-lab-code/BLI_processing_fitting), originally developed for BLI data analysis and used in ^37^.

### In silico de novo binder design with BindCraft

Binders for human Brd4^BD2^ were designed using BindCraft V1.5.2^27^, available at GitHub (https://github.com/martinpacesa/BindCraft). The software was executed in Google Colab using an NVIDIA A100 GPU and the human Brd4^BD2^ structure (Chain A in PDB: 5T35)^28^ was used as input. For BC001-BC012, no residues were targeted as hotspot residues and binder size was defined ranging from 40 to 140 residues. For BC021-BC038, residues 362, 365 and 404 were targeted as hotspot residues and binder size was set to range from 40 to 140 residues. BindCraft was configured with the following settings: design_protocol: default, prediction_protocol: default, interface_protocol: default, template_protocol: default and filter_option: default. A total of 40 designs were generated, ranked by decreasing interface predicted template modeling score (i_pTM). The 14 designs were selected based on the sequence diversity for experimental screening.

## Acknowledgements

We thank Prof. Kouhei Tsumoto and Dr. Ryo Matsunaga at The University of Tokyo for providing access to the BLI instrumentation through the AMED BINDS support platform. We also thank the technical staff Ms. Sakura Shimizu for DNA template preparation. Molecular graphics were made with UCSF ChimeraX, developed by the Resource for Biocomputing, Visualization, and Informatics at the University of California, San Francisco, with support from National Institutes of Health R01-GM129325 and the Office of Cyber Infrastructure and Computational Biology, National Institute of Allergy and Infectious Diseases.

## Abbreviations

AzPhe-CME: *p*-azido-L-phenylalanine cyanomethyl ester
BLI: Biolayer Interferometry
BioPhe-CME: *N*-biotinyl-L-phenylalanine cyanomethyl ester
eFx: Flexizyme
mNG: mNeonGreen
MetRS: methionyl-tRNA synthetase
ncAA: noncanonical amino acid
Sav: streptavidin

## Supplementary information

## Supplementary Methods

### Preparation of linear DNA templates for IVTT and *in vitro* transcription

Linear DNA templates for *in vitro* translation and *in vitro* transcription were generated by PCR using plasmids as templates and appropriate primers. Amplified DNA fragments were purified using the FastGene Gel/PCR Extraction Kit (FG-91302, NIPPON Genetics) and quantified with the Qubit dsDNA Quantification Assay Kit (Q32854, Thermo Fisher Scientific). All sequences used in this study are provided in Supplementary Tables S1–S7. Specifically, DNA oligonucleotide sequences are listed in Supplementary Table S1, amino acid sequences of BindCraft-designed binders targeting Brd4^BD2^ in Supplementary Table S2, amino acid sequences of individual proteins in Supplementary Table S3, nucleotide sequences of DNA templates for cell-free translation in Supplementary Table S4, DNA sequences of synthetic BD2 binder gene fragments used for fusion with mNG constructs in Supplementary Table S5, nucleotide sequences of DNA templates for in vitro transcription in Supplementary Table S6, and nucleotide sequences of tRNA^IniTx^ in Supplementary Table S7.

T7pro-XXX-mNG (all 64 start codons): The mNG fragment was amplified from a plasmid using primers Oligo01 and Olig02, followed by agarose gel electrophoresis and gel extraction purification. The purified DNA fragment was then used as a template for a second PCR to introduce start codon variants and append a T7 promoter sequence. This reaction employed three primers (Oligo02, Oligo03, and Oligo04–67), where Oligo04–67 contained all 64 possible start codon sequences. PCR was performed using KOD One polymerase in a 20 μL reaction containing 10 ng template DNA, 0.3 μM Olig02, 0.3 μM Olig03, and 3 nM Oligo04–67. The cycling conditions were as follows: 98 °C for 10 s; 5 cycles of 98 °C for 10 s and 74 °C for 10 s; 5 cycles of 98°C for 10 s and 72 °C for 10 s; 5 cycles of 98 °C for 10 s and 70 °C for 10 s; followed by 20 cycles of 98 °C for 10 s and 68 °C for 10 s.

T7pro-CAC-mNG Brd4^BD2^-targeting binder: The T7pro-CAC-mNG fragment was amplified from DNA template T7pro-CAC-mNG using Oligo68 and Oligo69 and purified by agarose gel electrophoresis followed by gel extraction. The BD2 binder fragment was synthesized as a gene fragment containing a 5′ overlap sequence for fusion to the mNG fragment and a 3′ sequence for downstream PCR amplification (see Supplementary Table S2 for sequences). The T7pro-CAC-mNG fragment and the BD2 binder fragment were fused by overlap extension and amplified by PCR using Oligo01 and Oligo68.

T7pro-mScarlet-Brd4^BD2^: The T7pro-mScarlet fragment was amplified from a plasmid using Oligo68 and Oligo70 and purified by agarose gel electrophoresis followed by gel extraction. The Brd4^BD2^ fragment was amplified from a linear DNA template (T7proN-SP6 RNAP-v4-BD2) described in the previous study^1^ using Oligo01 and Oligo71 and purified by gel extraction. The T7pro-mScarlet fragment and the Brd4^BD2^ fragment were fused by overlap extension and subsequently amplified by PCR using Oligo01 and Oligo68.

T7pro-tRNA^IniTx^ variants: Five tRNA templates, tRNA^IniTx02^_GGG_, tRNA^IniTx02^_GGA_, tRNA^IniTx02^_GUG_, tRNA^IniTx03^_GUG_, and tRNA^IniTx04^_GUG_ were prepared by primer extension followed by PCR amplification. First, each template was assembled by primer extension using the following oligonucleotide pairs: Oligo72/Oligo73 for tRNA^IniTx02^_GGG_, Oligo72/Oligo74 for tRNA^IniTx02^_GGA_, Oligo72/Oligo75 for tRNA^IniTx02^_GUG_, Oligo76/Oligo77 for tRNA^IniTx03^_GUG_, and Oligo72/Oligo78 for tRNA^IniTX04^_GUG_. Primer extension was performed with each primer at a final concentration of 1 μM. The cycling conditions were as follows: 98 °C for 10 s; 5 cycles of 55 °C for 10 s and 68 °C for 10 s. The resulting products were then used directly as templates for PCR amplification in a 40-fold scaled reaction containing 1× KOD One Master Mix and 1 μM each of Oligo79 and Oligo80. Oligo80 contained 2′-O-methyl modifications at its two 5′-terminal nucleotides to suppress non-templated nucleotide addition by T7 RNA polymerase^2^. The cycling conditions were as follows: 98 °C for 10 s; followed by 25 cycles of 55 °C for 10 s and 68 °C for 10 s.

T7pro-T-boxzyme: Two types of T-boxzymes with distinct sequences were prepared to enable aminoacylation by recognizing tRNA anticodons GTG and GGA. These were Tx2.1CAC, which recognizes the GTG anticodon, and Tx2.1UCC, which recognizes the GGA anticodon. DNA templates for transcription of T-boxzymes were generated using Oligo81/Oligo82 for Tx2.1CAC and Oligo83/Oligo84 for Tx2.1UCC, followed by PCR amplification using Oligo85/Oligo86. Primer extension was performed with each primer at a final concentration of 1 μM. The cycling conditions were as follows: 98 °C for 10 s; 5 cycles of 55 °C for 10 s and 68 °C for 10 s. The resulting products were then used directly as templates for PCR amplification in a 20-fold scaled reaction containing 1× KOD One Master Mix and 1 μM each of Oligo85 and Oligo86. The cycling conditions were as follows: 98 °C for 10 s; followed by 15 cycles of 55 °C for 10 s and 68 °C for 10 s.

## Supplementary Figures

**Figure S1.**
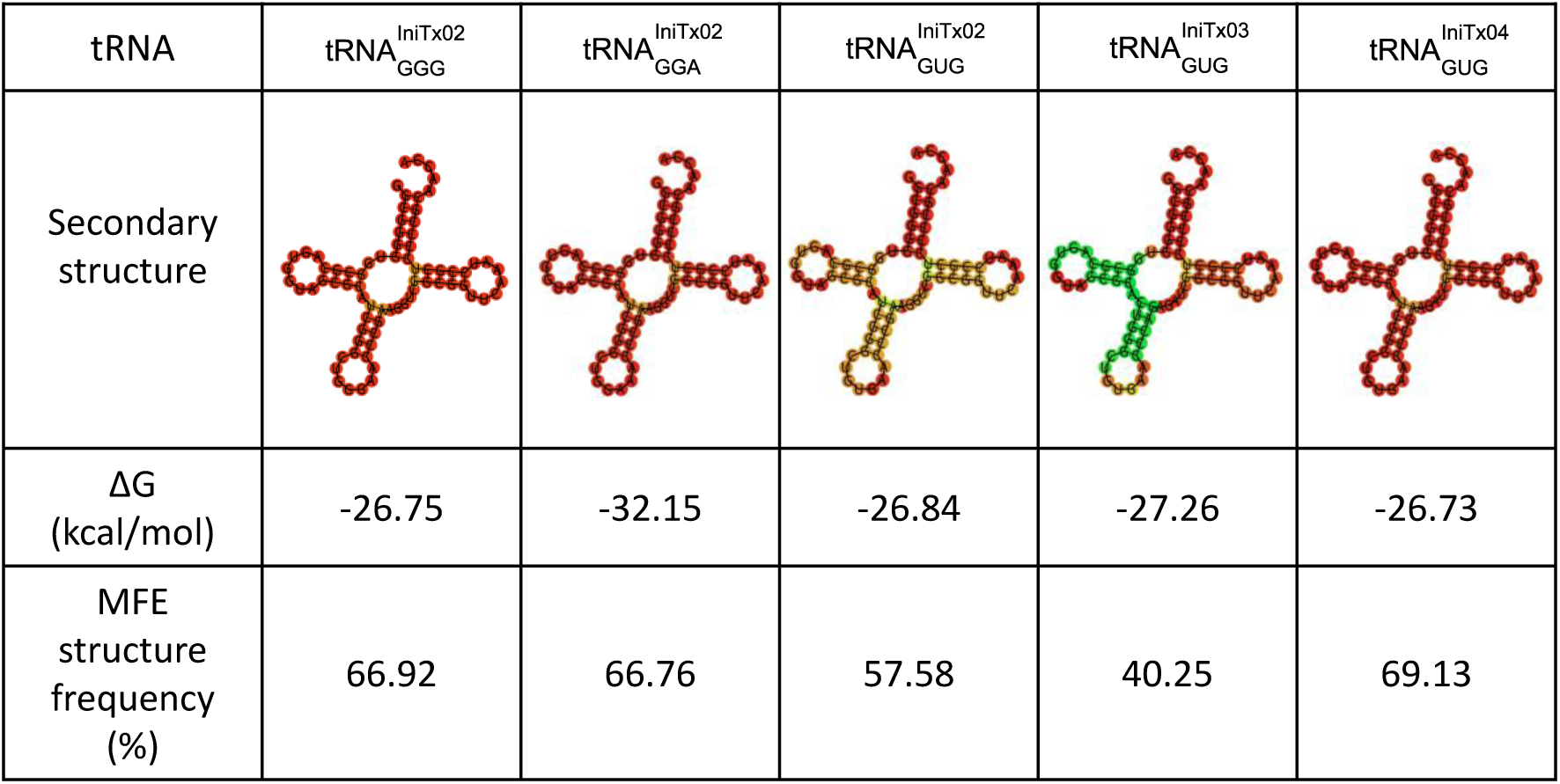
Predicted secondary structures and thermodynamic parameters of tRNA^IniTx^ variants. Secondary structures of the five designed tRNA^IniTx^ variants were predicted using RNAfold. Corresponding minimum free energy (MFE) values and predicted thermodynamic parameters are summarized alongside each structure.

**Figure S2.**
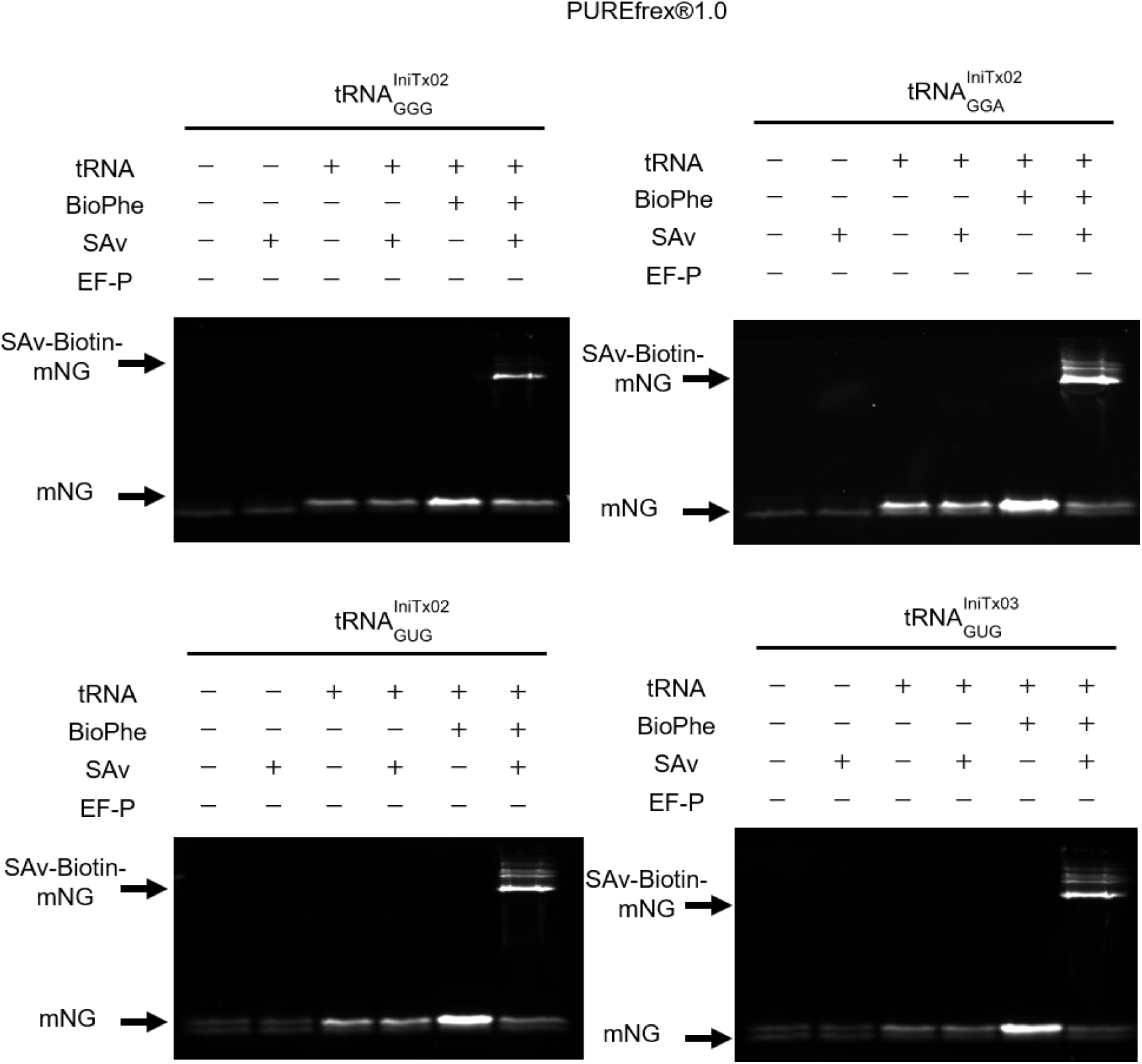
Detection of BioPhe incorporation into mNG in PUREfrex 1.0 without EF-P by SAv-dependent gel-shift assay. The gel was visualized based on the fluorescence of mNG in non-boiled SDS–PAGE.

**Figure S3.**
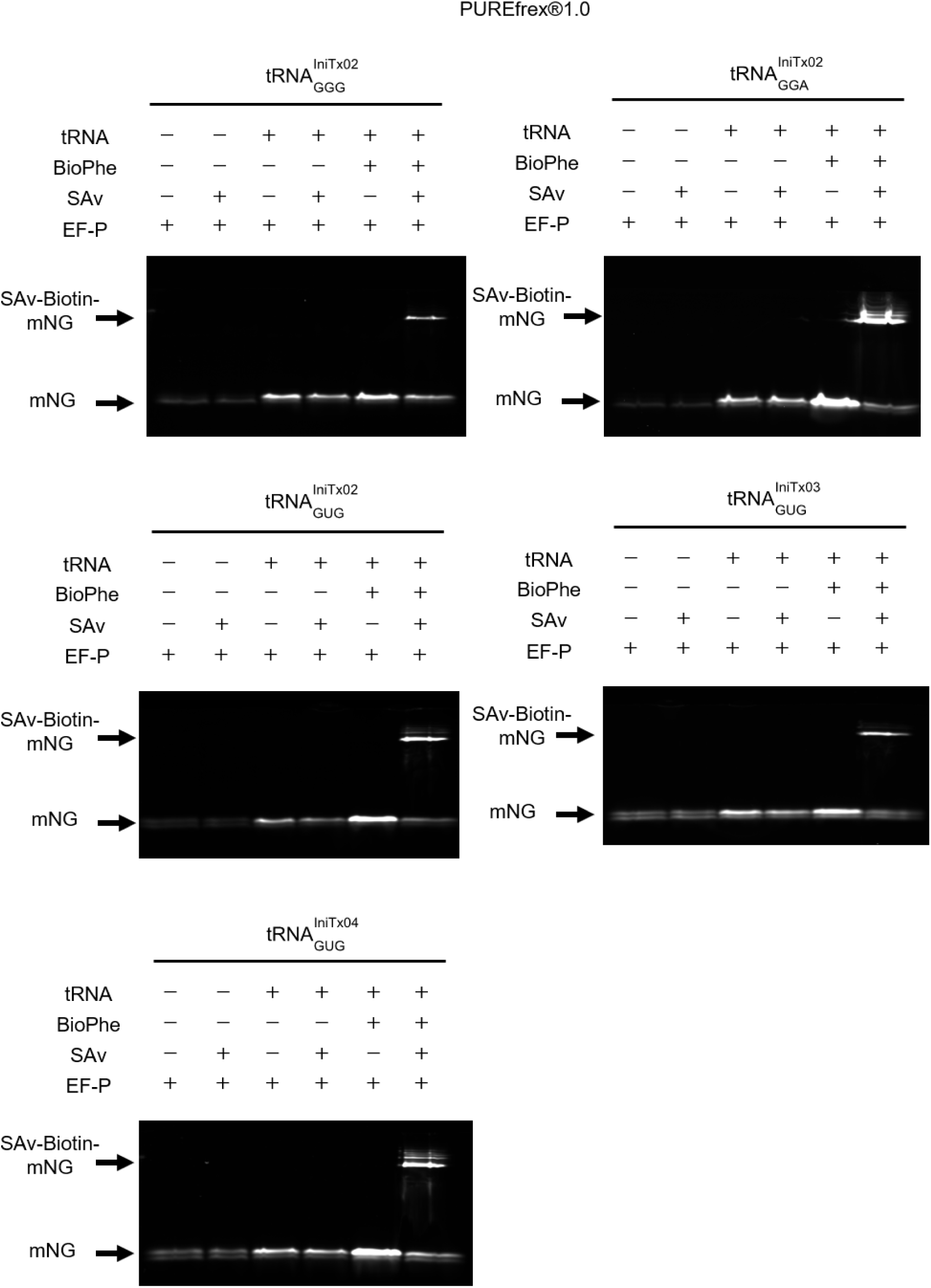
Detection of BioPhe incorporation into mNG in PUREfrex 1.0 with EF-P by SAv-dependent gel-shift assay. The gel was visualized based on the fluorescence of mNG in non-boiled SDS–PAGE.

**Figure S4.**
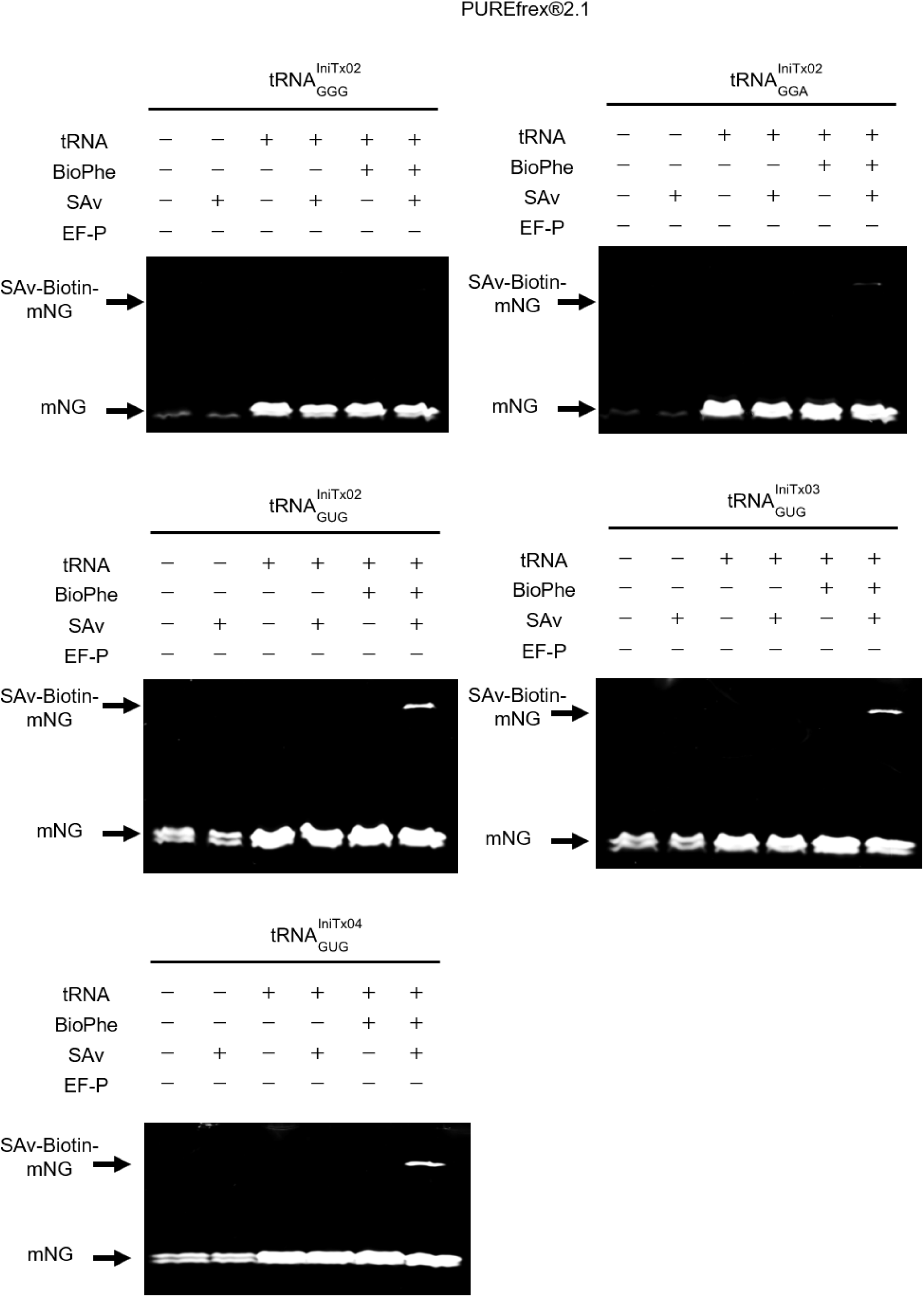
Detection of BioPhe incorporation into mNG in PUREfrex 2.1 without EF-P by SAv-dependent gel-shift assay. The gel was visualized based on the fluorescence of mNG in non-boiled SDS–PAGE.

**Figure S5.**
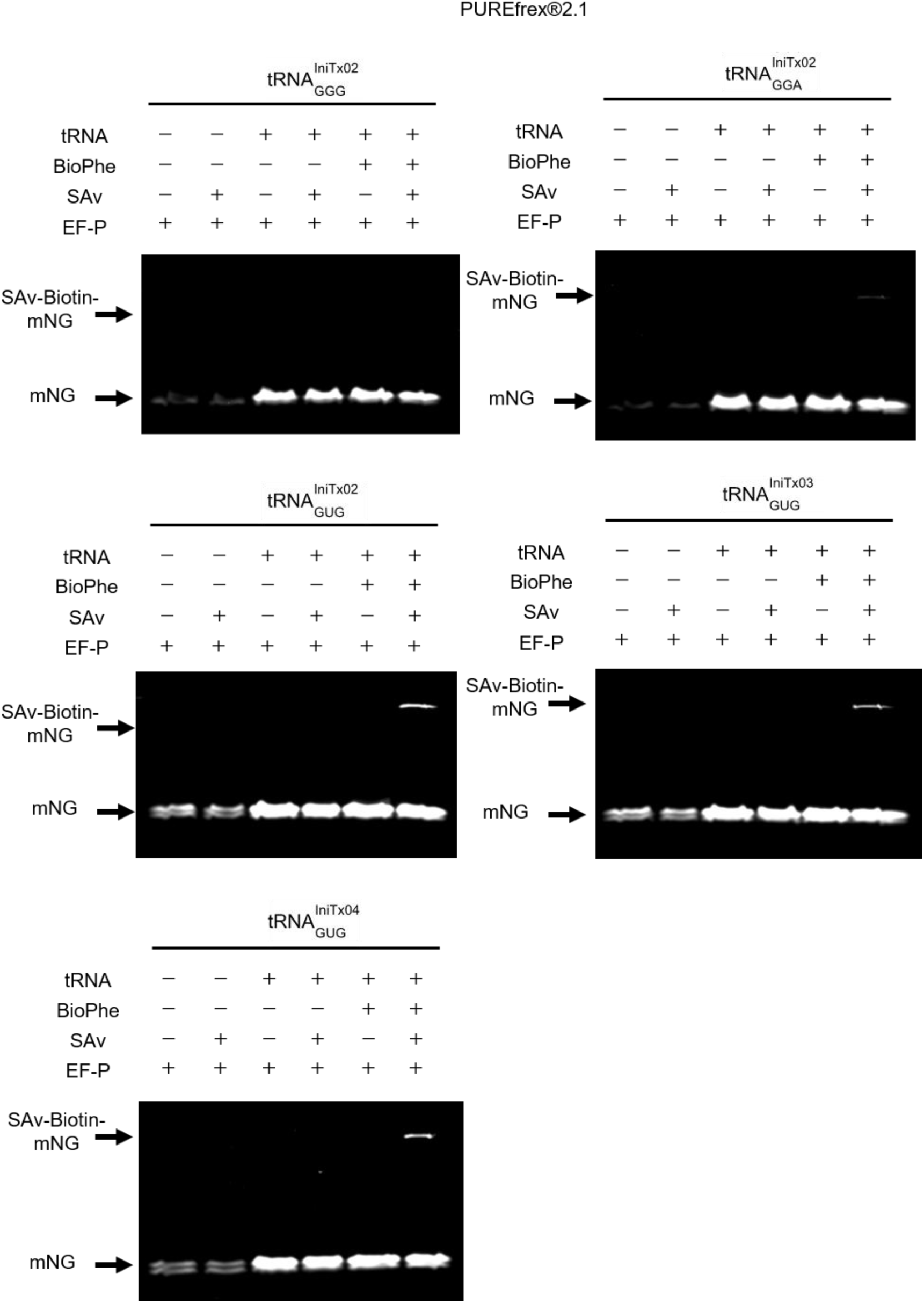
Detection of BioPhe incorporation into mNG in PUREfrex 2.1 with EF-P by SAv-dependent gel-shift assay. The gel was visualized based on the fluorescence of mNG in non-boiled SDS–PAGE.

**Figure S6.**
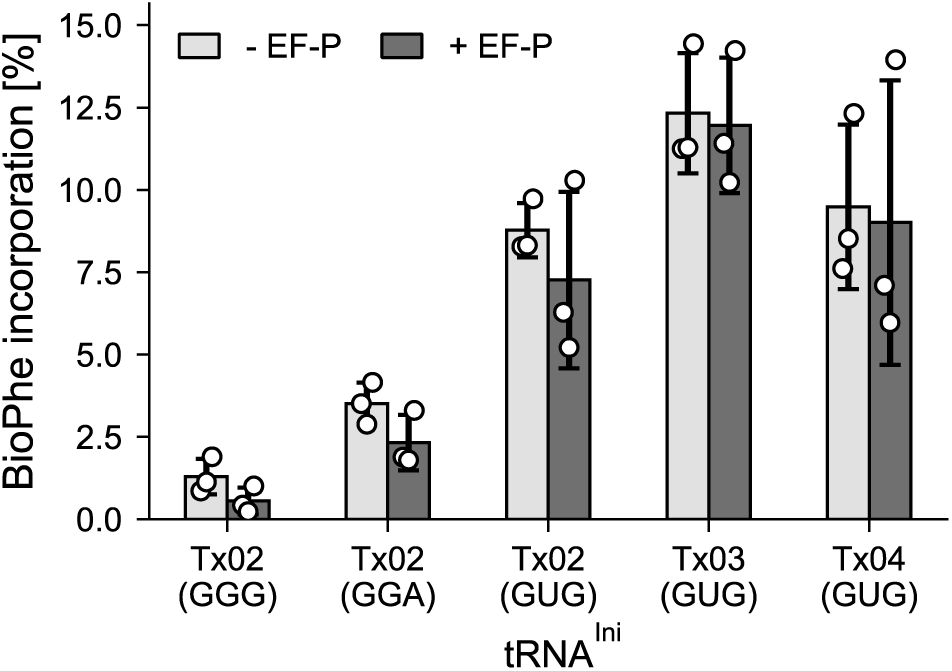
Comparison of BioPhe incorporation efficiencies among five tRNA^IniTx^ variants corresponding to artificial initiation codons in PUREfrex 2.1. The data represent the mean ± s.d. of three independent translation reactions (n = 3).

**Figure S7.**
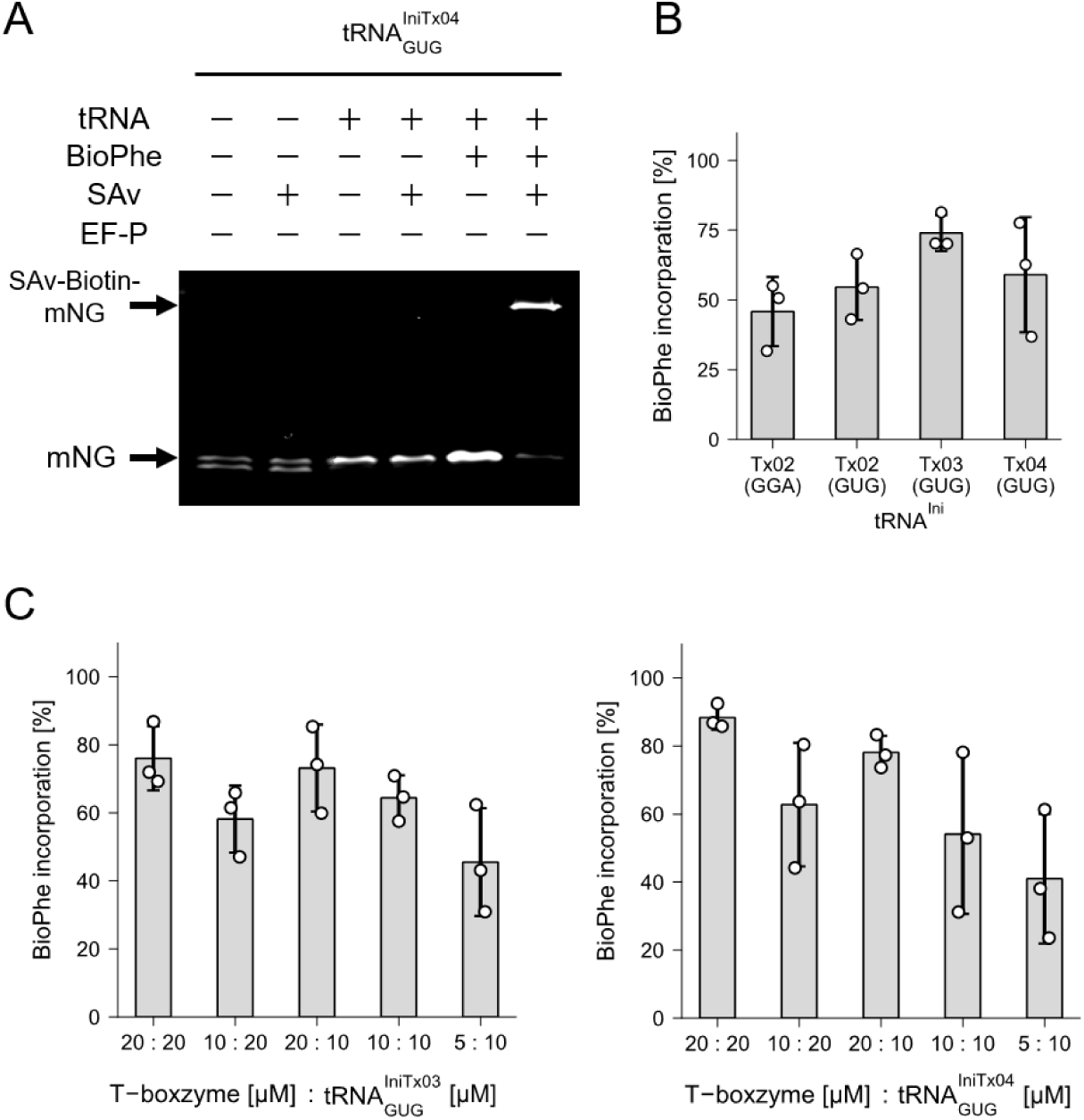
BioPhe incorporation by Tx2.1-mediated aminoacylation. (**A**) Detection of BioPhe incorporation using tRNA^IniTx04^_GUG_ into mNG by SAv gel-shift assay in non-boiled SDS–PAGE. (**B**) BioPhe incorporation efficiencies by four tRNA^IniTx^ variants cognate to selected artificial initiation codons. (**C**) Optimization of aminoacylation conditions by T-boxzyme for tRNA^IniTx03^_GUG_ (left panel) and tRNA^IniTx04^_GUG_ (right panel). Data in (B) and (C) represent the mean ± s.d. of three independent translation reactions (n = 3).

**Figure S8.**
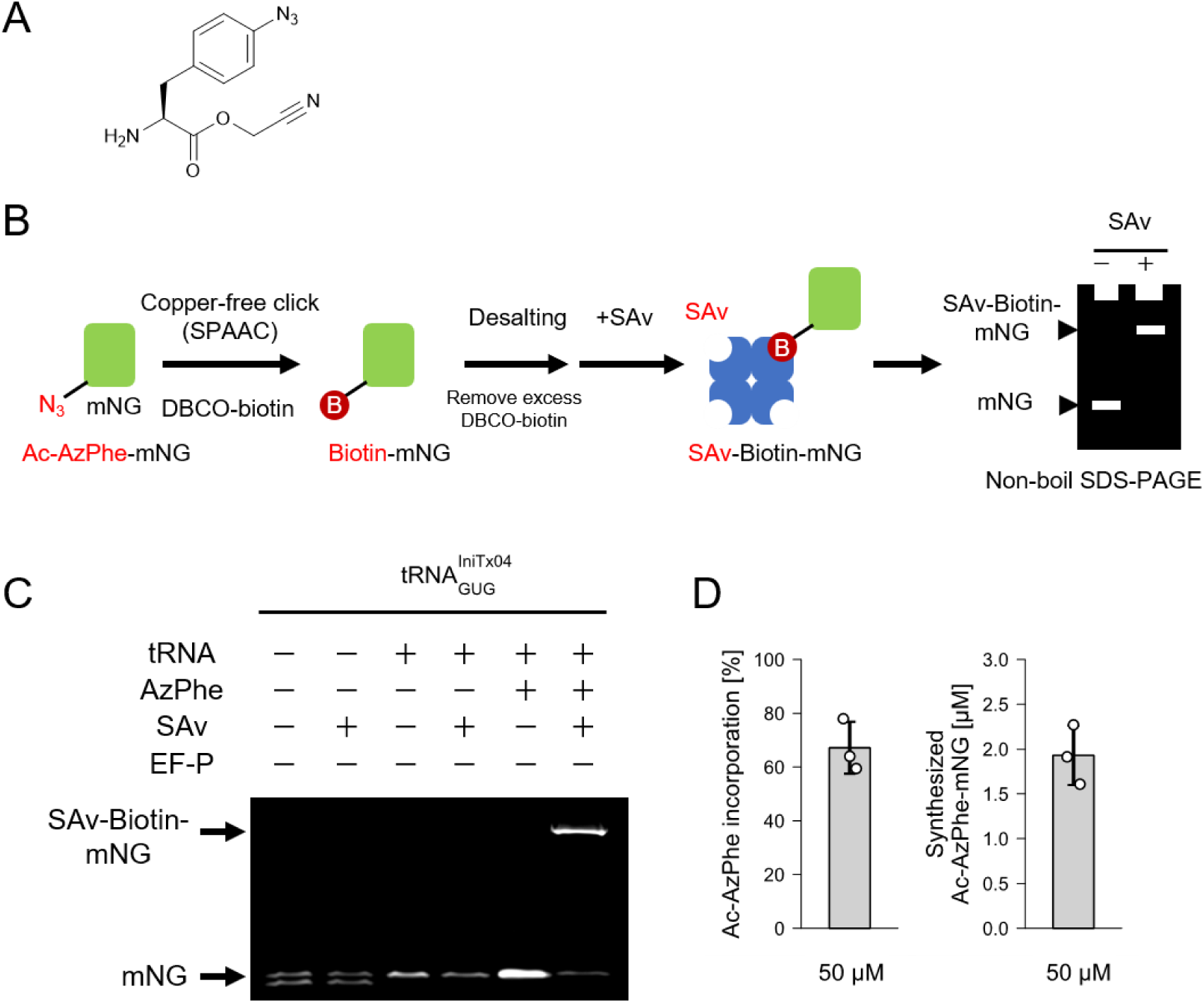
Evaluation of Ac-AzPhe incorporation using tRNA^IniTx04^_GUG_. (**A**) Chemical structure of AzPhe-CME. (**B**) Schematic representation of the workflow for detection of Ac-AzPhe-incorporated mNG. Ac-AzPhe-containing proteins were labeled with sulfo-DBCO-biotin via copper-free click chemistry, followed by desalting to remove unreacted DBCO-biotin. Biotinylated proteins were then incubated with streptavidin (SAv) and analyzed by non-boiled SDS–PAGE. (**C**) Detection of Ac-AzPhe incorporation using tRNA^IniTx04^_GUG_ into mNG by SAv gel-shift assay in non-boiled SDS–PAGE. (**D**) Quantification of Ac-AzPhe incorporation efficiency (left panel) and yield of Ac-AzPhe-mNG (right panel). Data represents the mean ± s.d. of three independent translation reactions (n = 3).

**Figure S9.**
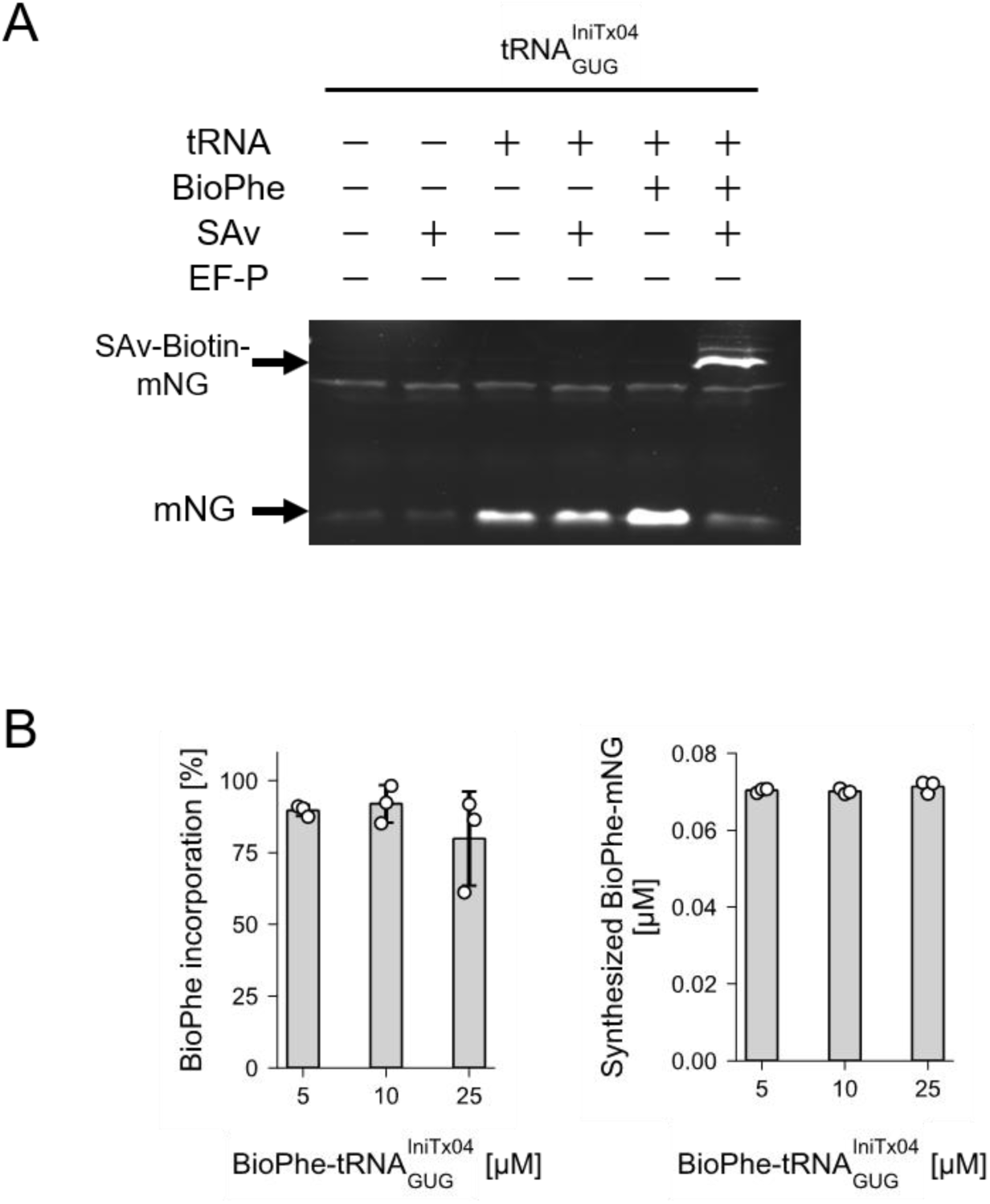
BioPhe incorporation in an *E. coli* extract–based cell-free translation system. (**A**) Detection of BioPhe incorporation using tRNA^IniTx04^_GUG_ into mNG by SAv gel-shift assay in non-boiled SDS–PAGE. (**B**) Effect of BioPhe-tRNA^IniTx04^_GUG_ concentration on incorporation efficiency (left panel) and absolute yield of BioPhe-incorporated mNG (right panel). Data represent the mean ± s.d. of three independent reactions (n = 3).

**Figure S10.**
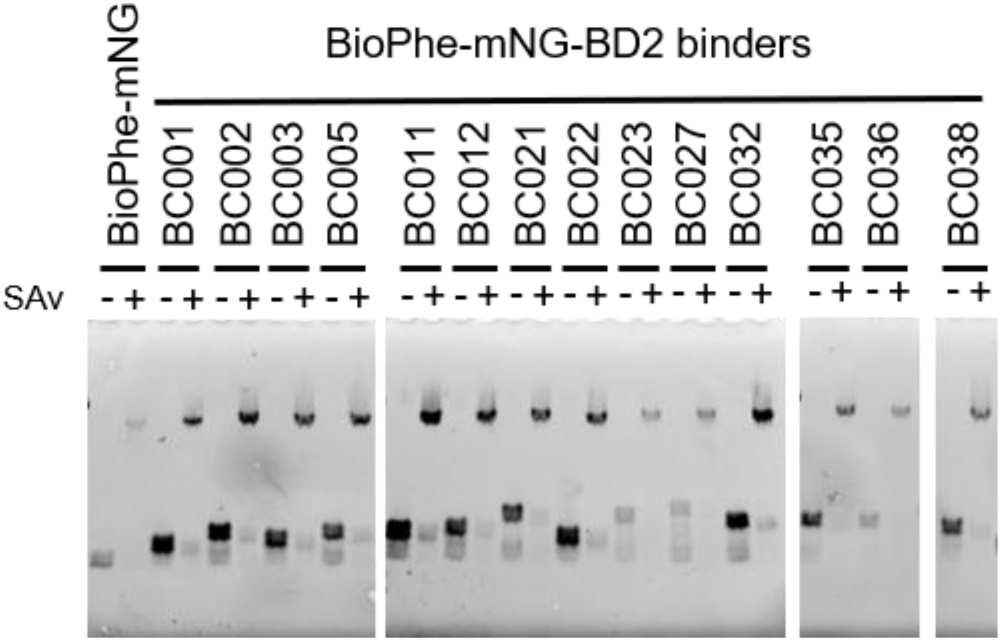
Expression analysis of BioPhe-mNG binders. Designed BioPhe-mNG-Brd4^BD2^ binders were expressed using PUREfrex 1.0 supplemented with BioPhe-tRNA^IniTx04^_GUG_. BioPhe incorporation into mNG was visualized by SAv gel-shift assay in non-boiled SDS–PAGE.

